# Gene mapping methodology powered by induced genome rearrangements

**DOI:** 10.1101/2022.05.11.491291

**Authors:** Hideyuki Yone, Hiromitsu Kono, Hayato Hirai, Kunihiro Ohta

## Abstract

Phenotypic variation occurs through genome rearrangements and mutations in certain responsible genes; however, systematic gene identification methodologies based on genome rearrangements have not been fully established. Here, we explored the loci responsible for the given phenotype using the TAQing system and compared it with a conventional mutagenesis-based method. Two yeast strains with different genetic backgrounds and flocculation phenotypes were fused and genomic rearrangements were induced by transient DNA breaks. Then, selection pressure was applied and multiple mutants were generated, showing different flocculation abilities. We also raised mutants with altered cohesiveness due to spontaneous mutations during long-term recursive passages of haploid strains without TAQing treatment. Comparative genomic analysis of the TAQed mutants revealed three chromosomal regions harboring pivotal flocculation genes, whereas conventional mutagenesis generated a more diverse list of candidate loci after prolonged selection. The combined use of these approaches will accelerate the identification of genes involved in complex phenotypes.

## Introduction

In traditional genetic mapping methods, genes involved in the expression of a particular phenotype are identified by generating many mutants of this phenotype through conventional mutagenesis. Since authentic mutagenesis often produces multiple mutations in the first generation of mutants, identifying the genes responsible for the phenotype requires further purification procedures, using backcrossing or verification procedures by reverse genetics to eliminate the effects of irrelevant mutations. Generally, this genetic analysis method works efficiently but requires a relatively long time for the alteration of generations during crossing. The conventional analysis of genotype-phenotype correlation using mutations in single genes is also gradually approaching saturation after considerable historical accumulation of similar trials.

In addition to these technical hurdles, certain known phenotypes are difficult to address using simple genetic analysis. For example, quantitative traits are altered to various extents by a combination of multiple genes with small effects. In the analysis of genes involved in such traits, quantitative trait loci (QTL) analysis, which statistically analyzes the linkage between genetic polymorphisms at multiple sites (genetic markers) and phenotypes after biparental mating, is used to clarify the complex involvement of multiple genes^1,2^. Another important approach is genome-wide association study (GWAS), which analyzes the association between the observed trait differences and the genome-wide nucleotide sequence variations of different individuals^3-5^. Moreover, expression QTL (eQTL) analysis, a correlation analysis method between transcript expression levels and genetic markers, is widely used in conjunction with GWAS to identify genes or alleles responsible for diseases and phenotypes, and databases and tools has become extensive^6-9^. However, these methods are based on the analysis of populations with diverse phenotypes and mutations that already exist or have been acquired through biparental mating or other means, thus limiting the range of samples for analysis.

Furthermore, phenotypic alterations occur through genome rearrangements such as copy number variations (CNVs), translocations (TLs), and loss of heterozygosity (LOH), as reported for phenotypes associated with cancer traits and various genetic diseases^10,11^ such as the Prader-Willi syndrome (heterozygous 15q11-q13 deletion)^12^, the Williams-Beuren syndrome (heterozygous 7q11.23 deletion)^13^, and the BCR/ABL1 translocation (t(9;22)(q34;q11))^14^. However, gene-mapping methods based on genome rearrangement have not been fully established. To resolve these technical difficulties, it is crucial to combine conventional mutation analysis with other methodologies based on genome rearrangements that do not involve time-consuming crossing or meiotic processes.

We previously developed the TAQing system technology that randomly induces multiple DNA double-strand breaks (DSBs) and subsequent recombination events by conditionally activating restriction endonucleases introduced in living yeasts and *Arabidopsis thaliana* mitotic cells^15,16^. In the original TAQing system, we introduced a gene encoding the thermo-activatable restriction enzyme TaqI, which recognizes the four-base TCGA sequence into living cells and is transiently expressed using inducible promoters. Upon elevating the temperature of the culture, TaqI in the cell is activated to randomly form DSBs throughout the genome, leading to a large-scale genome rearrangement by DNA repair. Importantly, the TAQing system generated point mutations at low levels. Additionally, genetic rearrangements induced by the TAQing system often generate remarkable phenotypic changes without genetic crossing and meiotic processes. These features enabled us to easily and quickly identify genes responsible for phenotypic changes.

We further extended the TAQing system to many non-conventional industrial yeasts, including *Candida utilis*, an imperfect fungus that cannot undergo normal meiosis and cannot be improved by crossbreeding. For this, we developed a TAQing system of the protein transfection type (TAQing2.0) by directly delivering TaqI into fungal cells using the cell-penetrating peptide method^17^. Additionally, we employed restriction endonucleases, other than TaqI (e.g., MseI) in the TAQing system, to improve the frequency of genome rearrangement and phenotypic diversification in plants (extended-TAQing system, Ex-TAQing)^16^.

Here, we developed a method to study the genotype-phenotype correlation by employing the TAQing system. We applied the TAQing system to cell-fused yeast strains with many single-nucleotide variations (SNVs) on each homologous chromosome and generated mutants with altered flocculation phenotypes by TAQing-induced genome rearrangements. Comparative genomic analysis of the mutants led us to identify loci that are important for the flocculation phenotype. This approach is faster than the conventional mutagenesis approach combined with reverse genetics. TAQing-based gene mapping is expected to facilitate genetic studies in various organisms.

## Results

### Selection of TAQed mutants with altered flocculation abilities

To investigate the genotype-phenotype correlation based on the TAQing system-inducing large-scale genome rearrangements, we focused on *Saccharomyces cerevisiae*’s flocculation phenotypes. Two yeast strains with different genetic backgrounds and flocculence phenotypes (S799/SK1 background and YPH499/S288C background) were fused and used for genetic analyses employing TAQing-induced genome rearrangements (Fig. 1a). The cell-fused diploid strain WT14 (*MATa/a*) had 0.7 % SNVs^18^, which enabled us to identify chromosomal rearrangement sites in the TAQed mutants. The WT14 strain showed much more cohesive phenotypes than both parental strains and readily precipitated under gravity (Fig. 1b, see WT14), possibly due to the trait’s enhancement by the complementation factors between the two parental S799 and YPH499 strains.

**Fig. 1.**
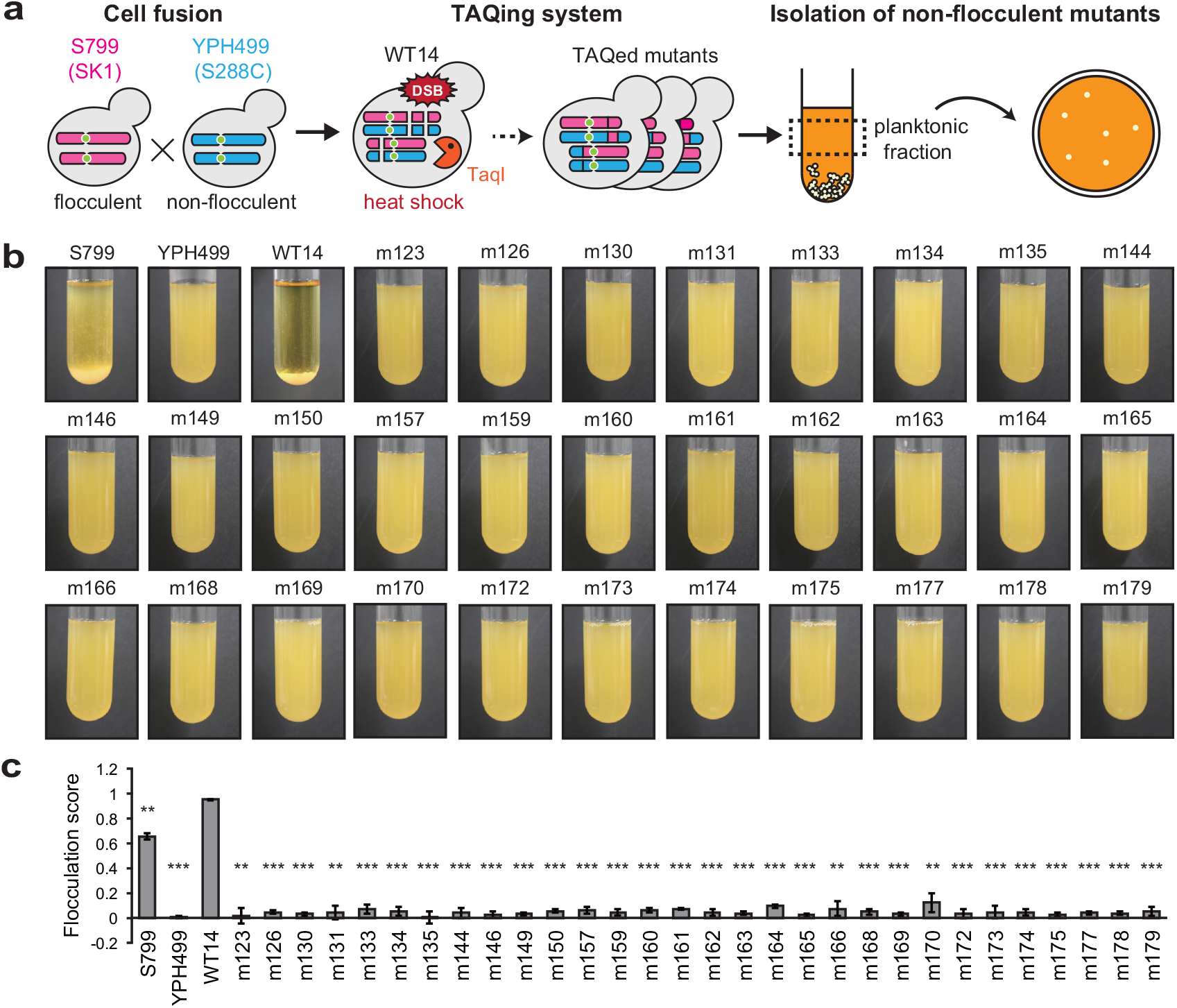
TAQing system alter flocculation phenotypes (a) A schematic of the workflow to obtain non-flocculent phenotypes from flocculent strains. *S. cerevisiae* haploid strains flocculent S799 and non-flocculent YPH499 are cell-fused into diploids (WT14), followed by TAQing treatment. Heat-activated endonuclease TaqI introduces multiple DNA double-strand breaks, leading to large-scale genome rearrangement. Non-flocculent TAQed mutants are isolated by screening planktonic cells with the remaining culture supernatant. (b) Flocculation behaviors of parental strains (S799, YPH499, WT14) and TAQed mutants (m123, m126, m130, m131, m133, m134, m135, m144, m146, m149, m150, m157, m159, m160, m161, m162, m163, m164, m165, m166, m168, m169, m170, m172, m173, m174, m175, m177, m178, and m179). (c) Flocculation scores of parental strains and TAQed mutants. *P* values were calculated using the Welch’s *t*-test compared to WT14: \*\**P* < 0.01 and \*\*\**P* < 0.001. Error bars represent standard deviation (*n* = 3).

We then induced genomic rearrangements in the fusion strain WT14 by TaqI-mediated DNA cleavage using the original TAQing system (Fig. 1a). The TAQed mutants were then subjected to a one-step procedure to select cells exhibiting reduced flocculation abilities from the cell population that remained in the upper part of the culture medium (planktonic fraction) (Fig. 1a). After inoculating the collected cells on agar plates, single colonies were isolated. We obtained 30 TAQed mutants exhibiting different levels of flocculation abilities (Fig. 1b), as revealed by the flocculation scores calculated from the observed cell sedimentation velocity of each strain (see more details in the Methods section, Fig. 1c). The parental S799 and fused WT14 strains showed very high flocculation scores, whereas the isolated TAQed mutants exhibited low flocculation scores despite some differences.

### Whole-genome sequencing of the TAQed mutants

Genomic sequences of these TAQed mutants were studied by short-read sequencing (>50 coverage; Illumina, San Diego, CA, USA), as described in the Methods section. We used SNVs between the S799 (SK1 background) and YPH499 (S288C background) strains to determine the regions with chromosomal rearrangements. The mapped sequences are illustrated in Fig. 2a and Supplementary Fig. 1 (blue and megenta segments represent sequences derived from the S799 and YPH499 strains, respectively). We observed multiple SNVs, break-induced repairs (BIRs), short gene conversions (sGCs), TLs, circularization, and aneuploidies in the TAQed mutants (see Supplementary Table 1 and examples of m126, m131, m159, m166, m174, and m177 in Fig. 2a, and summary of all BIRs, sGCs, TLs, and circularization in Fig. 2b). As previously reported^15^, two TL events occurred between the Ty transposable elements on chromosomes I and XVI (Fig. 2a, m131) and chromosomes I and III (Fig. 2a, m177). Interestingly, we detected self-circularization of one chromosome XV at the TaqI recognition sequence TCGA in *HXT11* and *NRT1* loci (Supplementary Fig. 2, m159).

**Fig. 2.**
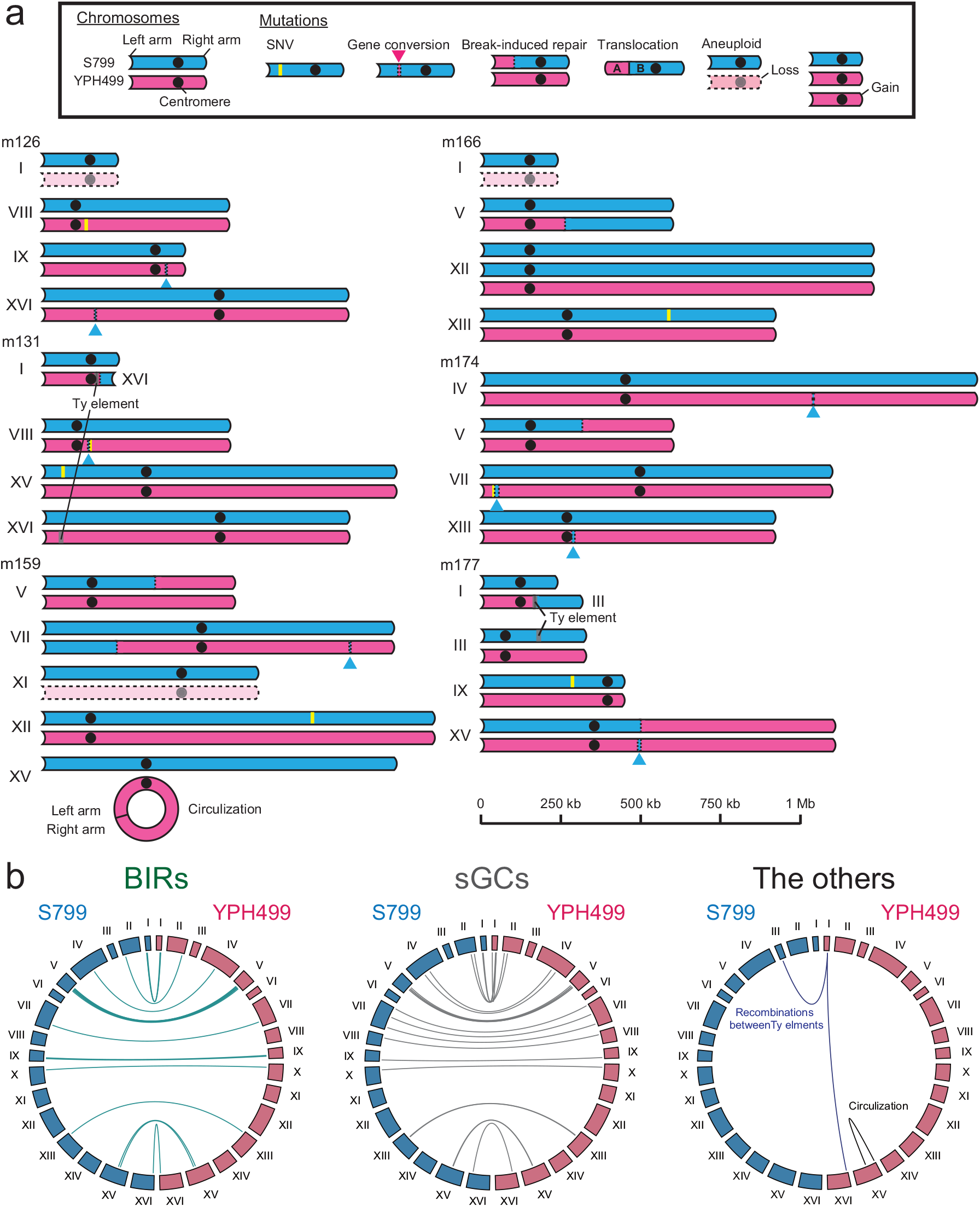
Chromosome structures in non-flocculent TAQed mutants (a) Schematic diagrams of rearranged chromosomes in non-flocculent TAQed mutants. S799 chromosomes, blue; YPH499 chromosomes, magenta. (b) Circular diagram of break-induced repairs (BIRs, green), short gene conversions (sGCs, gray), and the other rearrangements (purple, recombinations between Ty elements; black, circularization) within 30 non-flocculent TAQed mutants.

### Mapping of genes responsible for flocculation phenotypes

Notably, mutant 169 had a chromosomal rearrangement at the *FLO1* locus in the sub-telomeric region of chromosome I. In this strain, the region between *SWH1* and *FLO1* loci on YPH499-derived homologous chromosome I was replaced by the S799-derived counterpart, and the *FLO1* gene was recombined with the *TDA8* region in the left arm of chromosome I (Fig. 3a). *FLO1* is involved in the flocculation phenotype and encodes a cell wall lectin-like protein that can bind mannose^19^. The Flo1 protein (Flo1p) has a repetitive structure^20^ (Fig. 3b), and the length of its repeats varies from strain to strain^21^. Yeast strains with longer Flo1p repeat units exhibit higher flocculation ability^21^. Flo1p in S799 (SK1 background) and YPH499 (S288C background) had 10 and 18 repeat units, respectively (Fig. 3b, Supplementary Fig. 3), suggesting that YPH499 had a higher flocculation score than S799. However, we observed that the YPH499 haploid cells exhibited a very low flocculation score, and the WT14 strain (the hybrid of S799 and YPH499) had an even higher flocculation score than either parental strains (Fig. 1c).

**Fig. 3.**
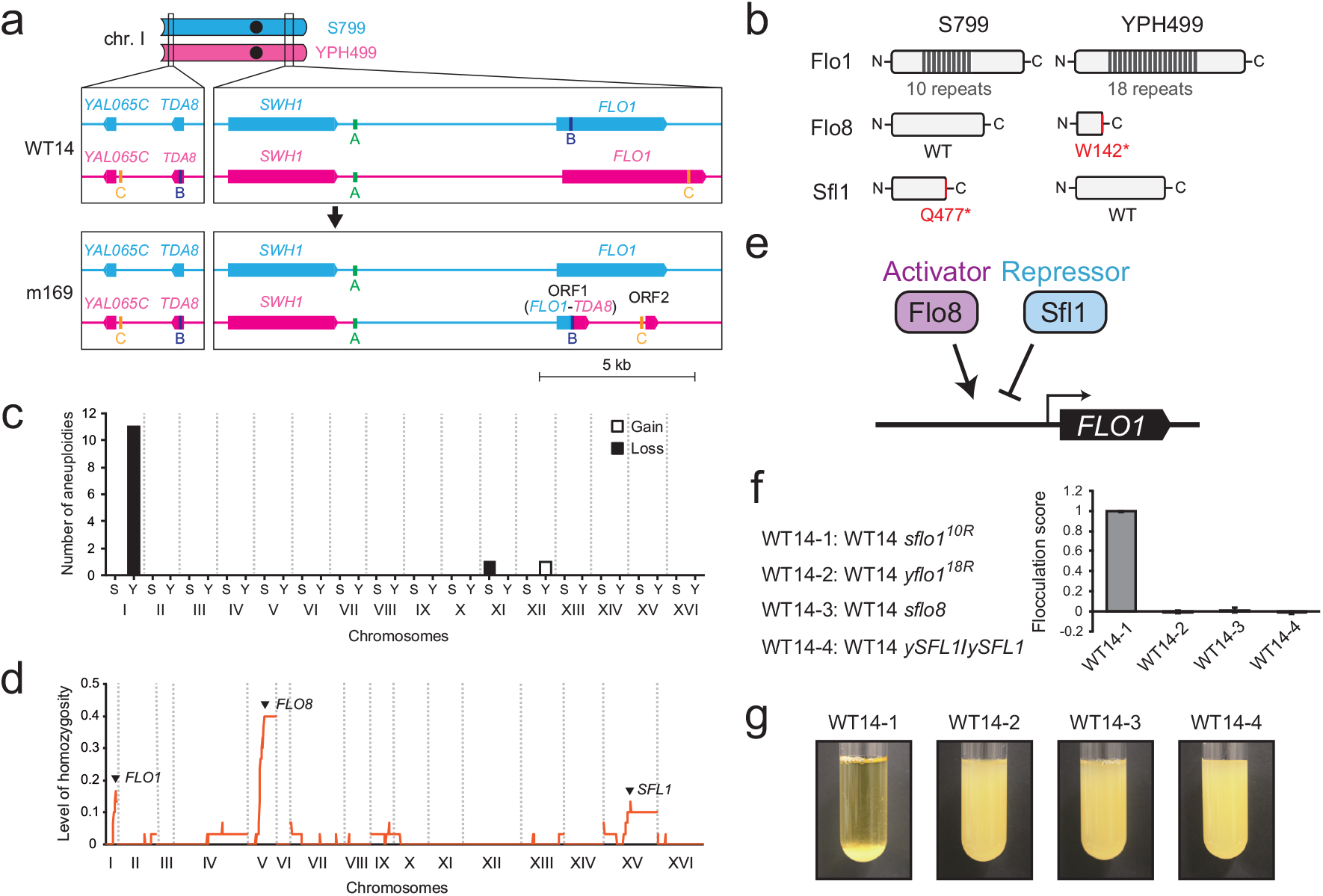
Comparative analysis of genomes of non-flocculent TAQed mutants (a) Complex gene conversions at subtelomeric regions in non-flocculent m169. Recombinations of homologous sequences A (green), B (navy blue), and C (yellow) lead to the depletion of the YPH499-derived *FLO1* gene and generates the two *de novo* ORFs. ORF1 is a fusion gene of *FLO1* and *TDA8* and ORF2 is the coding region of the *FLO1* 3’-terminus. (b) Natural variation in Flo1, Flo8, and Sfl1 proteins between parental strains. W142 in YPH499-derived Flo8 and Q477 in S799-derived Sfl1 were replaced by stop codons. (c) Number of aneuploidies within 30 non-flocculent TAQed mutants (S, S799; YPH499, Y). White bars show chromosomal gain and black bars show a chromosomal loss. (d) Level of homozygosity across the genome within 30 non-flocculent TAQed mutants, calculated as the ratio of homozygosity/heterozygosity in 10 kb windows. (e) Schematic diagram of the regulatory network of the *FLO1* gene. Flo8 activator and Sfl1 repressor have an antagonistic role in regulating the gene expression by binding the common promoter element. (f) Flocculation behaviors of WT14-1 (*sflo1*/*yFLO1*), WT14-2 (*sFLO1*/*yflo1*), WT14-3 (*sflo8*/*yflo8*), and WT14-4 (*ySFL1*/ *ySFL1*). (g) Flocculation scores of WT14-1 (*sflo1*/*yFLO1*), WT14-2 (*sFLO1*/*yflo1*), WT14-3 (*sflo8*/*yflo8*), and WT14-4 (*ySFL1*/ *ySFL1*). Error bars represent standard deviation (*n* = 3).

To solve this paradox, we analyzed the correlation between flocculation phenotypes and rearrangement events in multiple TAQed mutants with altered flocculation scores (Fig. 3c, d). Aneuploidy frequencies per chromosome revealed that TAQed mutants with reduced cohesion were more likely to lose chromosome I derived from YPH499 (Fig. 3c). Additionally, the analysis of LOH frequencies along each chromosome in the TAQed mutants revealed that LOH peaks (hotspots) were located in chromosomal regions containing flocculation-related genes *FLO1* (chromosome I), *FLO8* (chromosome V), and *SFL1* (chromosome XV) genes (Fig. 3d). These findings are intriguing because *FLO8* and *SFL1* encode transcription factors that either positively or negatively regulate *FLO1* gene expression, respectively^22-24^ (Fig. 3e). Moreover, the genomic sequences of the S799 and YPH499 strains suggested that *FLO8* in S799 and YPH499 strains are functional and nonfunctional (with a nonsense mutation at W142), respectively. A previous report also described that Flo1p is not expressed in S288C, because Flo8p in S288C has a nonsense mutation and is nonfunctional^25^ (Fig. 3b). Moreover, *SFL1* in S799 and YPH499 strains is likely nonfunctional (with a nonsense mutation at Q477) and functional, respectively^26^. These results suggest that the combination of polymorphisms in *FLO1, FLO8*, and *SFL1* loci via TAQing-induced genome rearrangements leads to various levels of reduced flocculation abilities.

To confirm this notion, we compared flocculation phenotypes of four strains using reverse genetics: WT14, S799 *FLO1* with 10 repeats (*sFLO1*^*10R*^)/YPH499 *FLO1* with 18 repeats (*yFLO1*^*18R*^), S799 *FLO8* (*sFLO8*)/YPH499 *flo8W142\** (*yflo8 W142\**), and S799 *sfl1Q477\** (*ssfl1Q477\**)/YPH499 *SFL1* (*ySFL1*); WT14-1, WT14, but deletion of *sFLO1*^*10R*^; WT14-2, WT14, but deletion of *yFLO1*^*18R*^; WT14-3, WT14, but deletion of *sFLO8*; WT14-4, WT14, but homozygous for *ySFL1* (Fig. 3f, g). Only WT14-1, in which *sFLO1*^*10R*^ was deleted in WT14, showed strong aggregation, while all other deletion strains showed severely reduced flocculation abilities. In other words, a significant reduction in flocculation ability was observed in the following three cases: 1) when *yFLO1*^*18R*^ (derived from YPH499, S288C) was deleted and only *sFLO1*^*10R*^ (derived from S799, SK1) was expressed; 2) when the functional wild-type *FLO8* (encoding a *FLO1* gene activator) was lost, or 3) when the functional wild-type *SFL1* (derived from YPH499, S288C, encoding a *FLO1* gene repressor) was homozygous in two copies. These results indicate that whole-genome sequencing of TAQed mutants with reduced flocculation abilities can efficiently enable us to identify a set of genes that play a central role in flocculation phenotypes.

### Gene mapping using a long-time selection with spontaneous mutagenesis

We then analyzed genotype-phenotype correlations using mutants with altered flocculation abilities generated by spontaneous mutagenesis during long-term passaging (Fig. 4a). In a similar experimental evolution, Hope et al. used haploid budding yeast cells to make genetic analysis easier^27^. They conducted long passages to generate cell lines with altered flocculation abilities and identify genes important for trait changes. Therefore, as the starting ancestor strain for the following iterative selections, similar to the analysis by Hope et al.^27^, we employed a YPH499-derived haploid strain (YPH499-*FLO8*) in which the wild-type *FLO8* derived from the S799 strain was expressed. Because the haploid YPH499-*FLO8* strain has both functional *FLO8* and *FLO1*^*18R*^ with longer repeats, it exhibits a very strong cohesive phenotype.

**Fig. 4.**
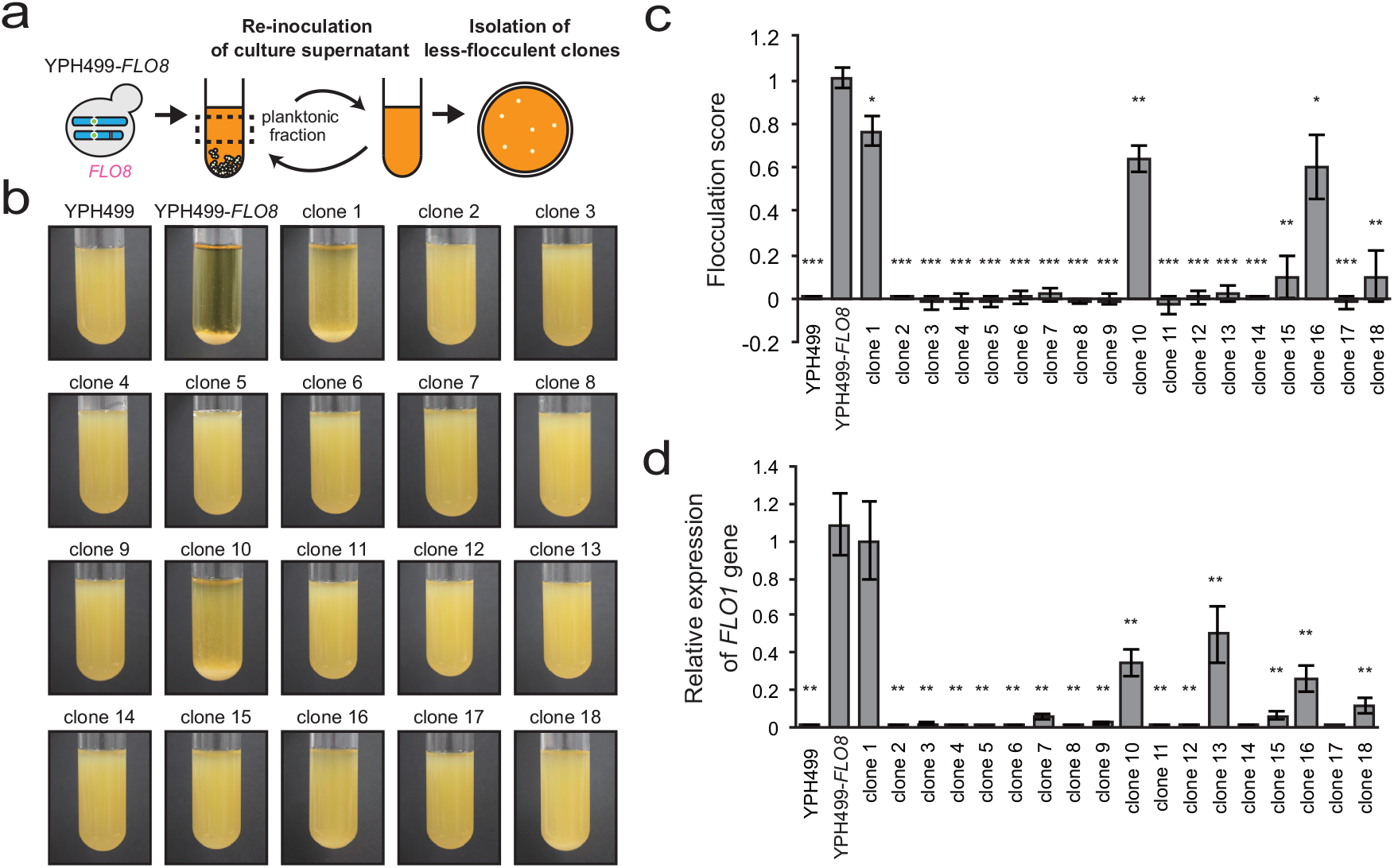
Altered flocculation phenotypes by re-inoculation of culture supernatant (a) A schematic of serial transfer experiment. YPH499 cells expressing the wild-type *FLO8* gene derived from the S799 strain (YPH499-*FLO8*) are cultured for 24 hours and culture supernatant was re-inoculated into a new medium. This manipulation was repeated for 20 days, followed by single colony formation on the agar plate. (b) Flocculation behaviors of YPH499, YPH499-*FLO8*, and less-flocculent clones isolated by serial transfer experiments. (c) Flocculation scores of YPH499, YPH499-*FLO8*, and less-flocculent clones. *P* values were calculated using the Welch’s *t*-test compared to YPH499-*FLO8*: \**P* < 0.05, \*\**P* < 0.01, and \*\*\**P* < 0.001. Error bars represent standard deviation (*n* = 3). (D) Relative expression of *FLO1* genes in YPH499, YPH499-*FLO8*, and less-flocculent clones. *FLO1* expression is measured by RT-qPCR and normalized to the expression of *ACT1* gene. *P* values were calculated using the Welch’s *t*-test compared to YPH499-*FLO8*: \*\**P* < 0.01, and \*\*\**P* < 0.001. Error bars represent standard deviation (*n* = 3).

We first cultured the ancestor strain YPH499-*FLO8* at 30 °C for 24 h with agitation in 18 independent test tubes containing a liquid medium. We temporarily stopped the agitation and withdrew the cells remaining in the planktonic fraction. The collected cells were diluted in a fresh liquid medium and allowed to grow for another 24 h, after which agitation was halted again and the planktonic fraction cells were collected. After repeating this procedure 20 times (20 days), we inoculated the final planktonic fraction cells onto agar plates and isolated single colonies from each of the 18 parallel experiments (Fig. 4a). The isolated cells were then grown in a liquid culture, and flocculation scores were calculated from the cell sedimentation rate of each strain (Fig. 4b, c). We obtained 18 mutant strains with various levels of reduced flocculation ability. Of these, partial reductions in flocculation abilities were observed in clones 1, 10, and 16, while other clones exhibited much more severe defects in cell-cell adhesion.

We then measured the expression levels of the *FLO1* gene, which encodes a cell wall protein involved in cell adhesion, in all 18 mutant strains (Fig. 4d). The results showed that the *FLO1* gene expression level was significantly decreased in almost all strains (17/18 strains), except for clone 1, which exhibited an almost identical score to that of the ancestral YPH499-*FLO8* strain. In clones 10, 13, and 16, a substantial amount of *FLO1* remained, although the expression level was decreased.

We determined the whole-genome sequences of these 18 mutant strains and compared them with the genomic sequence of the YPH499 strain to map the mutation sites (Table 1, Supplementary Fig. 4). We found a repeat length shortening (clone 1) or partial deletion (clone 13) in the *FLO1* gene, nonsense mutations in the *FLO8* gene (clones 4 and 17), and mutations in many other genes. The repeat length shortening of *FLO1* found in clone 1 was likely the reason for the reduced cohesiveness of this strain (Fig. 5a, Supplementary Fig. 5). A pop-out of a 15.9 kb region between *FLO1* and its 3’-side pseudogenic region caused the deletion in the *FLO1* gene found in clone 13 (Fig. 5a). Note that the alteration of *SFL1*, which was observed in the TAQing-based analysis, was not found in the mutation list. Moreover, the telomeric regions were deleted by over 10-20 kb in clones 5, 15, and 18. Clones 15 and 18 showed no gene deletions, whereas the *COS4* gene was deleted in clone 5 (Supplementary Fig. 4). Since the gene responsible for the flocculation phenotype was identified in clone 5, no analysis was performed on the *COS4* gene (see below).

**Table 1.**
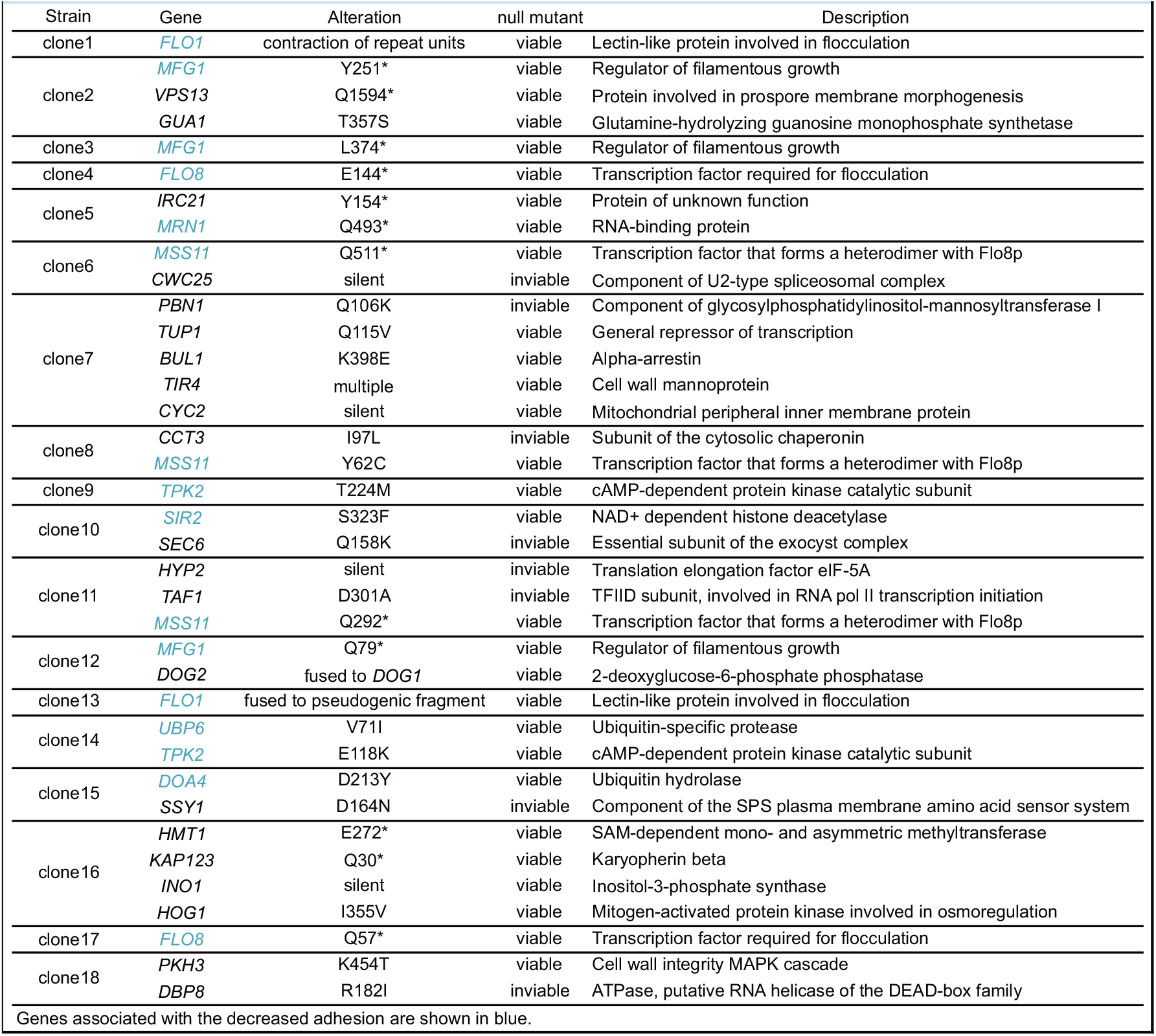
List of genes with de novo mutations in experimentally evolved clones.

**Fig. 5.**
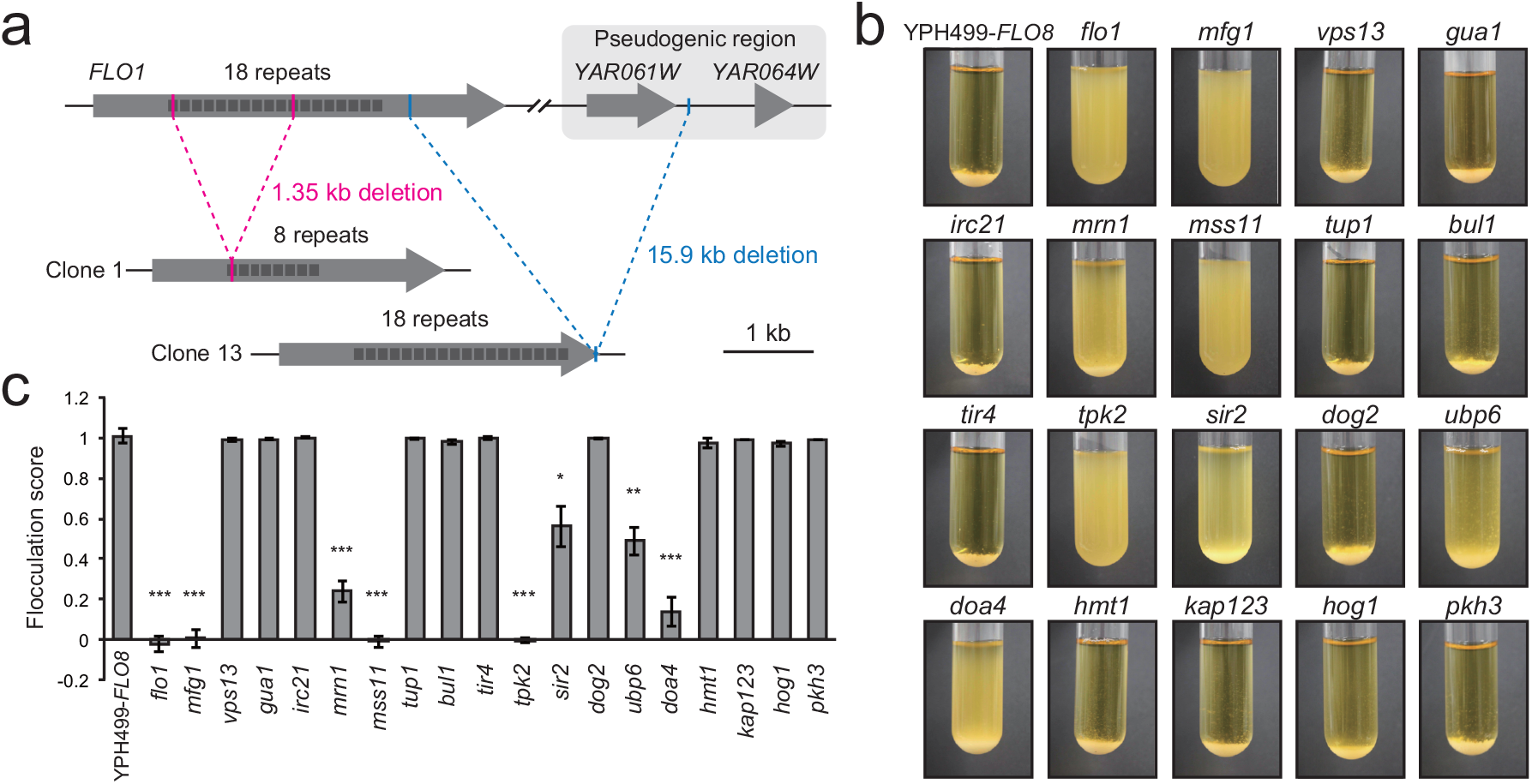
Genes mutated in less-flocculent isolates are associated with flocculation (a) Recombination of homologous sequences in the *FLO1* locus. Recombination between intragenic tandem repeats causes the contraction of the *FLO1* repeat (from 18 to 8 repeats) in clone 1. In clone 13, *FLO1* genes are fused to the homologous sequence in the pseudogenic region located about 16 kb downstream of the *FLO1* gene. (b) Flocculation behaviors of the deletion mutants of the YPH499-*FLO8* strain. (c) Flocculation scores of the deletion mutants. *P* values were calculated using the Welch’s *t*-test compared to YPH499-*FLO8*: \**P* < 0.05, \*\**P* < 0.01, and \*\*\**P* < 0.001. Error bars represent standard deviation (*n* = 3).

Based on the list of genetic mutations found among the 18 mutants, we examined their contribution to the altered flocculation phenotype by constructing individual gene disruptions (Fig. 5b, c). We detected markedly reduced flocculation scores in the single deletions of the *FLO1, MFG1, MRN1, MSS11, TPK2, SIR2, UBP6*, and *DOA4* genes. Fifteen of the 18 lines with reduced flocculation ability had mutations in one of these genes. Mfg1, Mss11, and Flo8 form transcription factor complex that induce the expression of downstream genes^28^. *TPK2* encodes cAMP-dependent protein kinase that inhibit transcriptional suppressor Sfl1^29^. Mrn1 is RNA-binding protein that repress the translation of target mRNAs including *SFL1*^30^. Sirtuin Sir2 is histone deacetylase required for the biofilm formation and flocculation in wine yeast strain^31^. *UBP6* and *DOA4* encode ubiquitin-specific proteases^32,33^, which have not been reported to be associated with flocculation to our knowledge. Clones 7, 16, and 18 harbored none of the studied mutations. In contrast, we observed no change in flocculation phenotypes in single deletions of *VPS13, GUA1, IRC21, TUP1, BUL1, TIR4, DOG2, HMT1, KAP123, HOG1*, and *PKH3*. We could not disrupt *CWC25, PBN1, CCT3, SEC6, HYP2, TAF1, SSY1*, and *DBP8* genes because they are essential. Therefore, the genes responsible for altered flocculation in clones 7 and 18 may be among these.

## Discussion

### Comparison of TAQing-based and mutagenesis-based genetic mapping methods

In this study, we proposed a method for genetic mapping using induced genome rearrangements in mitotic cells using the TAQing system. We compared the characteristics of this mapping method with those of conventional genetic mapping using spontaneous mutations in a model experiment for flocculation phenotypes. Large-scale genomic rearrangements induced by the TAQing system are triggered by a relatively clean DNA cleavage by restriction enzymes that can be repaired promptly. Therefore, unlike spontaneous mutations induced by replication errors or mutagenesis by chemical or radiation treatments, point mutations are rare, and phenotypic changes are caused mainly by genome rearrangements, including translocations, deletions, and copy number variation. This TAQing system feature enables easier identification of the important genes responsible for the trait of interest after whole-genome sequencing and comparative genomics studies.

Determining the whole-genome sequences of a large number of mutants and executing comparative genomic analysis is extremely costly and laborious. However, recent advances in DNA sequencing technologies have enabled rapid and low-cost genome sequencing of a large number of individuals and cells, thereby overcoming the technical challenges of TAQing-based genetic mapping. In fact, in this study, genome resequencing of approximately 33 strains, including the original tester strains, was conducted with more acceptable costs and time compared to previous studies. After genome resequencing, the chromosomal regions involved in the altered flocculation abilities could be efficiently identified using analytical tools for detecting aneuploidy and LOH in Fig. 2a, b. Accumulated information from gene annotation and gene ontology also strongly supports gene identification using the TAQing system. After analyzing the genomic sequences of numerous TAQed mutants, we could narrow down potential chromosomal regions involved in the phenotypic changes and further focus on candidate genes that may be involved in the phenotype by gene ontology analysis. Additionally, more rapid and automated analyses will be available if we refine the gene-mapping tools for bioinformatics analysis.

In this study, using TAQing-based mapping, we identified a group of genes involved in flocculation phenotype changes. We observed an extreme bias in the incidence of aneuploid formation on chromosome I in TAQed mutants with reduced flocculation abilities (we could detect only the loss of YPH499/S288C-derived chromosomes). Additionally, the frequency of LOH occurrence peaked in chromosome regions harboring *FLO1, FLO8*, and *SFL1* genes. The *FLO1* gene is present on chromosome I and is reported to encode a mannose-binding cell wall protein with intramolecular repeats, whose length correlates with flocculation strength. Many TAQed mutants (11 / 30 cases) with reduced flocculence lacked the YPH499/S288C-derived chromosome I, which contains the *FLO1* gene with a longer repeat. These TAQed mutants have only the S799/SK1-derived chromosome I harboring the *FLO1* gene with a shorter repeat, thereby exhibiting reduced cohesiveness.

It is noteworthy that in this experiment, along with the *FLO1* polymorphism, LOHs were highly concentrated in chromosomal regions, including *FLO8* and *SFL1*, which encode transcription factors that positively and negatively regulate *FLO1* gene expression, respectively. The *FLO8* gene encodes a transcription factor that activates the *FLO1* gene, and the *SFL1* gene product represses *FLO1* expression, thus suppressing flocculation phenotypes. Although natural yeasts often exhibit high flocculence, yeast strains with high flocculation abilities are difficult to handle in laboratories, and flocculence has been artificially reduced over many years of experimental processes (laboratory domestication). The laboratory strain YPH499, with the S288C background, lacks the functional *FLO8* gene and has low flocculence^25^. In contrast, the S799 strain derived from the SK1 strain, which is substantially distant from the S288C strain, retained a functional *FLO8* gene. Moreover, the S799 strain with the SK1 background contains a nonsense mutation in the *SFL1* gene that suppresses the high flocculation phenotype, whereas YPH499 with S288C contains the wild-type *SFL1* gene^26^. Thus, it is likely that TAQing-induced genome rearrangements altered the combination of these genetic variants, resulting in the diversification of flocculation phenotypes.

The fact that TAQing-based gene mapping could selectively shed light on three pivotal genes (*FLO1, FLO8*, and *SFL1*) involved in the flocculation phenotypes suggests that the TAQing system is indeed advantageous for the efficient identification of genes responsible for the phenotype of interest. However, for TAQing-based gene mapping, it is desirable to arrange the original tester multiploid strain with a considerable number of genetic polymorphisms (SNVs) along the homologous chromosomes. However, even if no interchromosomal polymorphisms exist in the tester strain, where parental chromosomes cannot be distinguished by SNVs (e.g., homozygous pure lines), we can still identify the genes responsible for the altered traits by analyzing CNVs and TLs occurring in the TAQed mutants.

In contrast, conventional gene identification using spontaneous mutagenesis can be effective even in the absence of SNVs on homologous chromosomes. In our analysis using spontaneous mutagenesis, we identified candidate genes, even in haploid cells with no homologous chromosomes. However, in mapping using spontaneous mutagenesis, multiple mutations are often introduced simultaneously. Therefore, to identify the most critical mutation leading to phenotypic changes, it is necessary to rule out the effects of irrelevant mutations by conducting additional backcrossing or reverse genetics experiments, which are laborious and time-consuming. In this study, TAQing-based mapping enabled us to analyze the phenotype-genotype correlation after only a single selection cycle, but the analysis using spontaneous mutagenesis required 20 iterations of the selection cycles, which was approximately 20 times as long as that in TAQing-based mapping.

While only three critical genes were identified as candidates in the TAQing-based method, over 20 candidate genes, including *FLO1*, were listed in the mapping using natural mutagenesis. This feature may be a major advantage of mapping using natural mutagenesis. It can list a wider range of responsible genes that could not be detected in the TAQing-based mapping, although mapping by natural mutagenesis requires tremendous additional effort and time to verify whether they are indeed involved in the flocculation phenotypes by disrupting the genes one by one.

### Relationship with QTL and GWAS

QTL and GWAS are powerful conventional methods for genetic mapping. QTL mapping analyzes quantitative trait gene regions by identifying known genetic markers that exhibit statistically significant linkages with trait differences in F2 offspring constructed by the mating of parents with different levels of quantitative traits. QTL mapping identifies gene regions based on the genetic diversity obtained from a single meiotic division after mating with a few parents. Therefore, the analysis requires time for genetic crosses and has a scale limitation.

In contrast, mapping by GWAS uses the genetic diversity of existing populations of related species with many genetic backgrounds derived from historical meiotic recombination events. Therefore, there is no need to perform time-consuming crossings to analyze the correlation between DNA markers and traits. Moreover, once the sequence information is obtained, we need only time for the data analysis. Although GWAS is generally considered more convenient than QTL mapping, there is a risk of listing false-positive candidates when dealing with rare traits, linkage disequilibrium, or peculiarities in the population used for analysis. In contrast, QTLs using biparental mating can sometimes enable the mapping of related genes that are difficult to identify in GWAS.

The TAQing system involves a precise comparison of the whole-genome sequences of the parental strains and the TAQed mutants with mitotically-induced genome rearrangements (CNVs, SNVs, TLs, and LOHs) that are linked to phenotypic alterations. Thus, the TAQing system employed an efficient GWAS-like mapping strategy using artificially acquired genetic diversity in mitotic cells. In other words, TAQing-based mapping combines the advantages of both GWAS and QTL methods and can be applied to a wider range of trait analyses in a short period and at low cost. TAQing-based mapping also applies to species without appropriate genetic markers or SNVs. In addition, since meiotic recombination is not required for TAQing-based mapping, it can be applied to interspecific hybrids that cannot form offspring and to hybrid cells by cell fusions, thus greatly expanding the target of genetic mapping.

### Comparison with existing mapping using mitotic cells

There are other mapping methods using mitotic cells in which chromosomes are recombined by ionizing irradiation^34^ or site-specific recombination mediated by CRISPR/Cas9^35^. Ionizing irradiation induces numerous point mutations owing to the generation of DNA chemical adducts formed by free radicals in cells. The presence of numerous irrelevant point mutations can hinder rapid gene identification and often forces us to conduct further verification processes, such as backcrossing or reverse genetics experiments. The CRISPR-based mapping method enables efficient identification of the genes responsible for trait changes by analyzing LOH and CNV, as in TAQing-based mapping. However, the preparation and transfection many gRNA for CRISPR/Cas9 target sites throughout the genome incur substantial costs. Such technical limitations do not exist in TAQing-based mapping, as there are numerous four-base cutter target sites in the genome where DNA breaks occur randomly. Laureau et al. achieved efficient genetic mapping in sterile yeast using yeast return-to-growth experiments^36^. However, there is a technical challenge with this method, which requires transient activation of meiotic recombination and cannot be applied to cells with intrinsic defects in meiotic recombinases.

As described above, the gene mapping method using the TAQing system proposed in this study is not only advantageous for identifying important genes involved in complex phenotypes, but also saves time and labor by using mitotically induced large-scale genome rearrangements instead of meiotic recombination events. They can also be applied to sterile hybrids or somatic cell genetics. This method, in combination with conventional mapping using mutagenesis, is expected to expand the possibility of identifying useful, unidentified genes involved in complex phenotypes.

## Methods

### Yeast strains, culture, and mutagenesis-based selection

The yeast strains YPH499 (S288C-derived haploid; *MATa ura3-52 lys2-801 ade2-101 trpl-Δ63 his3-Δ200 leu2-Δ1*) and S799 (SK1-derived haploid; *MATa ura3 lys2 ho::LYS2 leu2Δ arg4-bgl cyh2-z*)^37^ were used as parental strains of cell-fused strain WT14. Thirty TAQed mutants, namely m123, m126, m130, m131, m133, m134, m135, m144, m146, m149, m150, m157, m159, m160, m161, m162, m163, m164, m165, m166, m168, m169, m170, m172, m173, m174, m175, m177, m178, and m179 were isolated by TaqI activation in WT14 followed by the screening of non-flocculent cell populations. The hyperflocculent YPH499 strain (YPH499-*FLO8*) expressing the wild-type *FLO8* gene derived from the S799 strain was used as the ancestral strain in mutagenesis-based iterative selection. The naturally mutagenized clones 1-18 were generated by 20 cycles of selection as follows: 10 µL of planktonic fractions (supernatants of cultures after standing for 1 min) of the ancestral YPH499-*FLO8* cells were withdrawn and inoculated into 10 mL of fresh yeast extract-peptone-dextrose-adenine (YPDA) medium in 18 test tubes, followed by the culture at 30 °C for 24 h. Yeast cells were also cultured in SD/monosodium glutamic acid (MSG) medium at 30 °C for subsequent experiments.

### Yeast strain construction

Gene disruption was performed according to conventional yeast gene-targeting methods^38,39^. WT14-1, WT14-2, WT14-3, and W14-4 were constructed by fusion of YPH499 with S799 carrying *flo1*, S799 with YPH499 carrying *flo1*, S799 carrying *flo8* with YPH499, and S799 carrying *ySFL1* (wild-type *SFL1*) with YPH499, respectively. S799 carrying *ySFL1* was constructed by replacing the *ySFL1* cassette with the *sfl1::URA3* allele of the S799-derivative *sfl1* disruptant, followed by the selection of URA^-^ colonies on SD medium plates containing uracil and 5-fluoro-orotic acid (5-FOA, FUJIFILM Wako Pure Chemical Corporation, Japan). The YPH499-*FLO8* strain for natural mutagenesis experiments was constructed by integrating the *sFLO8:URA3* cassette into the intergenic region between *YCRdelta11* and *FEN2* on chromosome III in YPH499.

Cell fusion was performed using the following method, with some modifications to a previous study^40^. The parental strains grown to the log phase were harvested and washed once with sterile water. The cell pellets were suspended in a protoplasting solution (2.8 mg/mL Zymolyase -20 T (07663-91, Nacalai Tesque, Japan), 1.7 % 2-mercaptoethanol (Nacalai Tesque, Japan), and 25 µL/mL glusulase (PerkinElmer, Inc., USA) in MP buffer) and incubated at room temperature for 30 min. The protoplast cells were harvested and washed twice with MP buffer (1 M sorbitol, 100 mM NaCl, and 10 mM acetic acid, pH 5.5). Two parental protoplasted cells were mixed with 2 mL of 60 % polyethylene glycol and 0.2 mL of calcium chloride and incubated for 3 min. Next, 6 mL of MP buffer was added to the suspension, and the mixture was incubated for 6 min. Protoplast cells were washed twice with MP buffer, then added to the regeneration agar medium (0.67 % yeast nitrogen base without amino acids, 2 % glucose, 1 M sorbitol, 3 % agar, and CSM-his-trp-arg), and seeded into 90 mm dishes. Only diploid cells formed colonies on the agar plate, which enabled the selection of appropriate strains.

### TAQing system and screening of non-flocculent TAQed mutants

The WT14 cells harboring the TaqI expression vector were cultured in 10 mL of SD/MSG medium at 30 °C. The medium was supplemented with 150 µM CuSO_4_ and cultured for 4 h to induce TaqI expression as described by Muramoto et al^15^. Cells were washed in distilled water, incubated in a YPDA medium for 30 min at 42 °C to activate TaqI, and cultured for 24 h at 30 °C. After 24 h, the cell culture was agitated with a mixer for 1 min and allowed to stand for 1 min. Ten microliters of the supernatant (planktonic fraction) were plated on YPDA agar plates for single colony isolation. We measured the flocculation scores of the isolated TAQed mutants, as described below.

### DNA preparation for whole-genome sequencing

Yeast genomic DNA was extracted using the Dr. GenTLE™ High Recovery for Yeast Kit (Takara Bio Inc., Japan), according to the manufacturer’s instructions. The extracted genomic DNA’s quality was confirmed by agarose gel electrophoresis, and the concentration was measured using a Qubit dsDNA HS assay kit (Qubit 3, Thermo Fisher Scientific Inc., USA). Genomic DNA was shared to a size of approximately 300-400 bp by a Covaris Focused-ultrasonicator M220 (Covaris, LLC., USA). DNA libraries were prepared using the NEBNext Ultra II DNA Library Prep Kit for Illumina (New England Biolabs, USA) and NEBNext Multiplex Oligos for Illumina (New England Biolabs, USA). The quality of the DNA libraries was confirmed by electrophoresis using MultiNA (Shimadzu Corporation, Japan) as described by Muramoto et al^15^. DNA libraries were sequenced using Illumina HiSeq X Ten (> 50 coverage; Illumina, Inc., USA).

### Next generation sequencing data analysis

The sequencing reads (.fastq) were mapped using a Burrows-Wheeler Alingner to the reference genome sequences obtained from previous studies^15,41^. Small variants such as SNVs and deletions were called by Freebayes^42^ and filtered by vcffilter in vcftools (DP> 15 & MQM> 30 & QUAL/AO > 10 & SAF> 0 & SAR> 0 & RPR> 1 & RPL> 1 & QUAL> 100 & AF = 1) and bcftools^43^. LOHs and aneuploid chromosomes were determined by the ratio of coverage using the Integrative Genomic Viewer (IGV)^44^. Structural variations were detected by extracting chimeric sequence reads and searching for rearranged positions using SAM tools^45^ as described by Muramoto et al^15^. Circular plots visualizing genome-rearrangement events in Fig. 2b were generated by OMGenomics Circa 1.2.2 (https://omgenomics.com/circa/). PCR primer used to confirm chromosomal circularization in m159 were 5′-CCAAGCGATACCAGGTAGACCGGGAG-3′ and 5′-CTGGCACACCTTCCAGCCACATAGT-3′.

### Quantification of flocculation phenotypes

Cells were cultured in 10 mL YPDA medium for 24 h at 30 °C and suspended in a mixer for 1 min, followed by incubation in the culture tubes for 1 min. The flocculating cells precipitated at the bottom of the culture tubes, while planktonic cells remained in the suspension. The planktonic population (200 µL) was collected, and the optical density at 600 nm (OD600) was measured using iMark™ Microplate Reader (Bio-Rad Laboratories, Inc., USA). Clumped cells in the remaining culture were dispersed by adding 10 mM EDTA and the OD600 of the cell suspension was measured. The flocculation score (the ratio of flocculating cells to total cells in the culture) was calculated using the following formula:

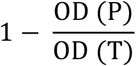

OD (P) and OD (T) are the OD600 of the culture supernatant and cell-dispersed culture, respectively.

### RNA extraction and RT-qPCR

RNA extraction and reverse transcriptase-quantitative polymerase chain reaction (RT-qPCR) experiments were performed according to the methods described by Hirota et al.^46^ and Galipon et al.^47^ with some modifications. Briefly, after being cultured in 10 mL of YPDA medium for 24 h, the cells were centrifuged at 3,000 × g for 5 min and frozen in liquid nitrogen. Frozen cell pellets were resuspended in 250 µL bead buffer (75 mM NH_4_OAc, 10 mM EDTA, pH 8.0) at 65 °C with 200 µL acid-washed glass beads (Sigma-Aldrich, USA), 25 µL of 10 % SDS, and 300 µL of acid-phenol: chloroform (pH 4.5, Thermo Fisher Scientific Inc., USA). The samples were stirred three times 1 min each at 1 min intervals and incubated at 65 °C, followed by a 10 min incubation at 65 °C, 1 min stirring, and 15 min centrifugation at room temperature (16,000 × g). The aqueous phase was transferred to a fresh tube containing 200 µL of bead buffer and 400 µL of phenol/chloroform/isoamyl alcohol (25:24:1, Sigma-Aldrich, USA). Tubes were stirred briefly and centrifuged at 4 °C, 16,000 × g for 15 min. The aqueous phase was transferred to a fresh tube with 600 µL isopropanol and 20.4 µL of 7.5 M ammonium acetate. Tubes were stirred briefly and centrifuged at 4 °C, 16,000 × g for 30 min. After discarding the supernatant, the pellet was washed with 70 % ethanol, air-dried, and resuspended in RNase-free water.

The PrimeScriptRT Reagent Kit with a gDNA eraser (Takara Bio Inc., Japan) was used to eliminate genomic DNAs and reverse transcription according to the manufacturer’s instructions. Real-time PCR was performed using the StepOne Real-Time PCR system (Thermo Fisher Scientific Inc., USA) and KAPA SYBR FAST qPCR Master Mix (2X) Kit (Sigma-Aldrich, USA). RT-qPCR primers for the *FLO1* gene were 5′-CGCCGATCACATCAACGAACT-3′ and 5′-ACCCCATGGCTTGATACCGTC-3′, and those for the *ACT1* gene are 5′-CTCCACCACTGCTGAAAGAGAA-3′ and 5′-CCAAGGCGACGTAACATAGTTTT-3′. *FLO1* expression was normalized to that of the *ACT1* gene.

## Acknowledgements

This work was supported by the AMED Grant Number JP20wm0325003, the Japan Science and Technology Agency (JST) CREST JPMJCR18S3 to K. O, and JST SPRING-GX Grant Number JPMJSP2108 to H. Y. We would like to thank Editage (www.editage.com) for English language editing and Miki Tamura for next generation sequencing library preparation.

## Author contributions

H. Y., H. K., H. H., and K. O. conceived the study; H. Y. and H. K. designed and performed genomic analyses and mutagenesis experiments. H. Y. and H. K. determined the whole-genome sequences of the yeast strains. All experiments were performed under the supervision of K. O.; K. O., H. Y., and H. K. wrote the manuscript, and all authors read and commented on the manuscript.

## Competing financial interests

The authors declare no competing financial interests but the following competing non-financial interests:

**Supplementary Fig. 1.**
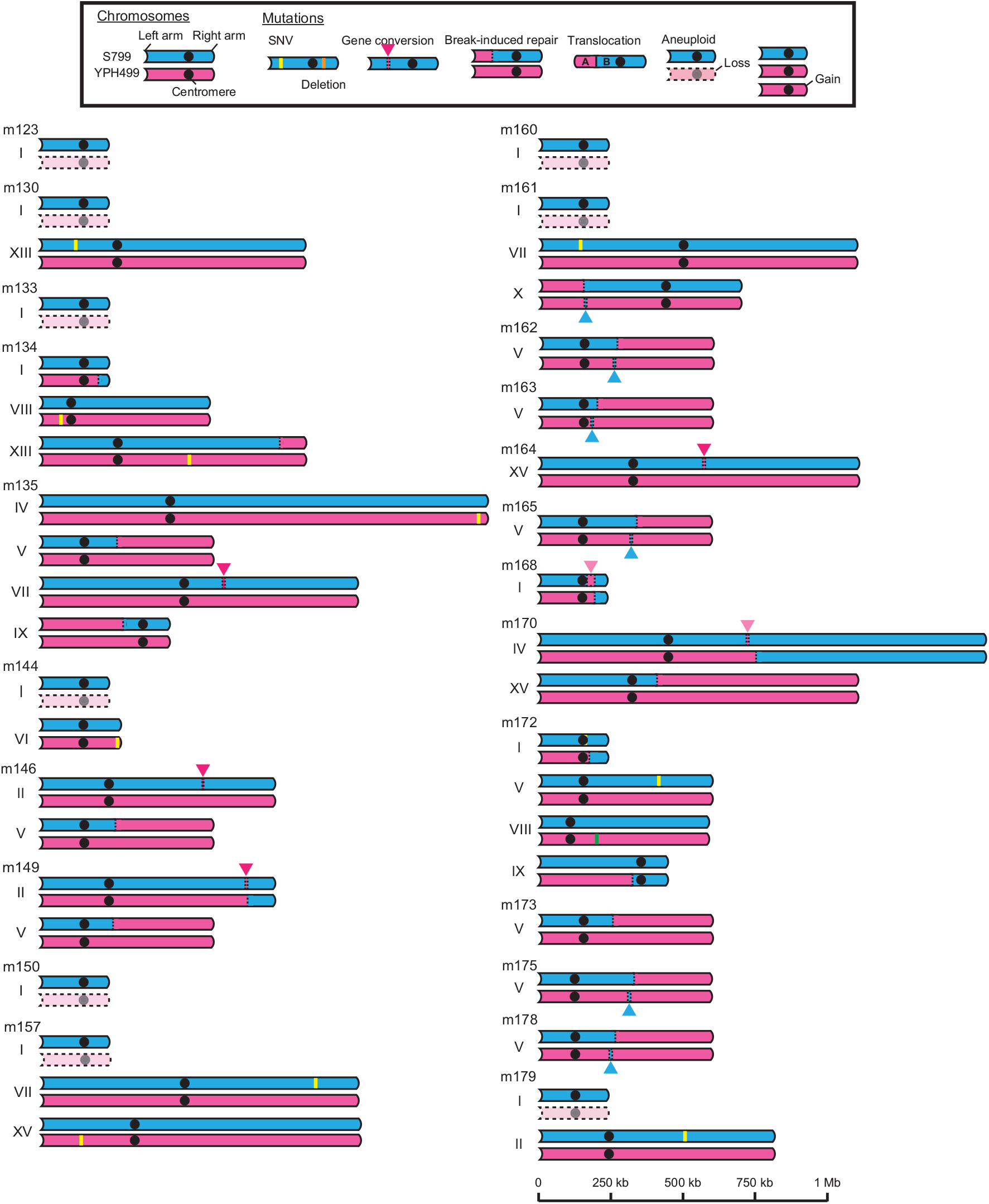
Rearranged chromosomes in TAQed mutants. Schematic diagrams of rearranged chromosomes in non-flocculent TAQed mutants m123, m130, m133, m134, m135, m144, m146, m149, m150, m157, m160, m161, m162, m163, m164, m165, m168, m170, m172, m173, m175, m178, and m179. S799 chromosomes, blue; YPH499 chromosomes, magenta.

**Supplementary Fig. 2.**
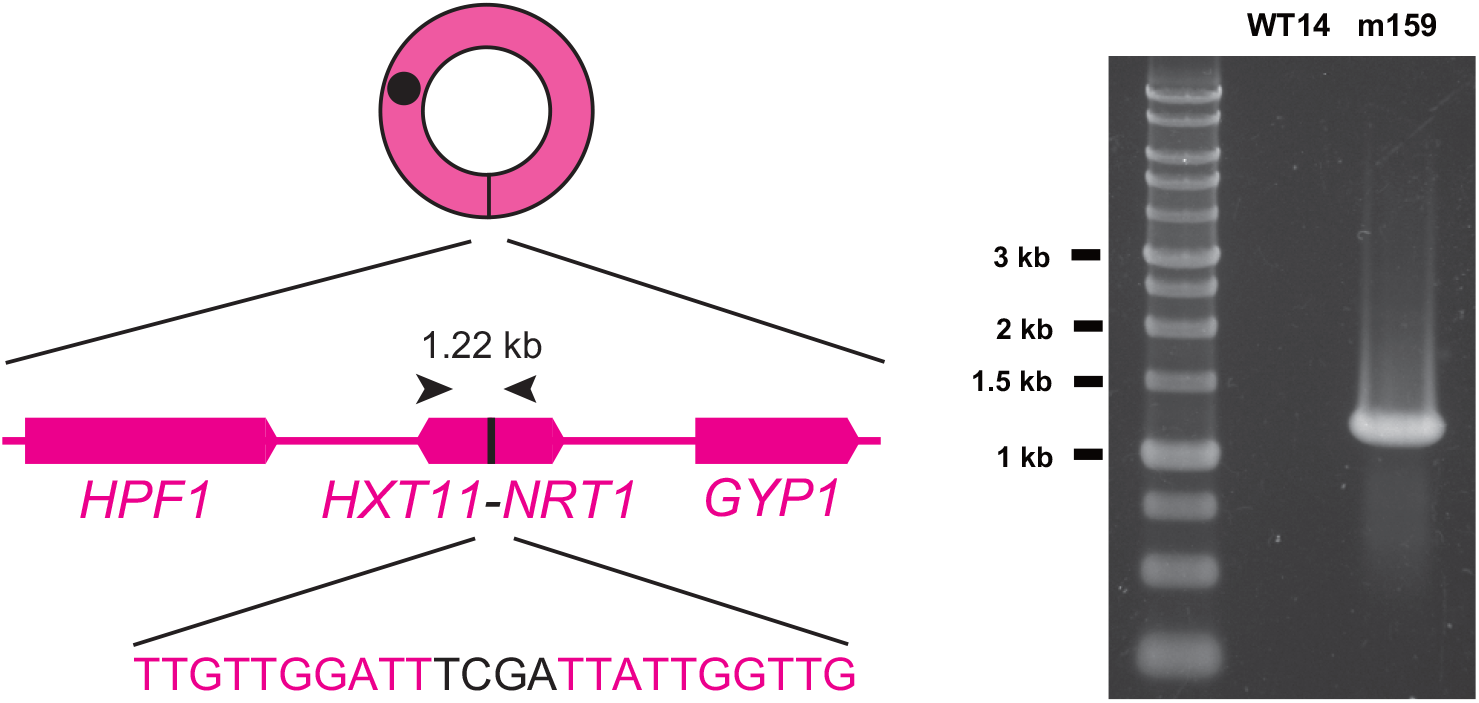
Breakpoint sequence of circular chromosome in m159. Breakpoints are at TaqI-recognition sequences (5′-TCGA-3′) located within *HXT11* and *NRT1* gene in YPH499-derived chromosome XV. Chromosomal circularization is verified by polymerase chain reaction (PCR). PCR primers are indicated by black arrows.

**Supplementary Fig. 3.**
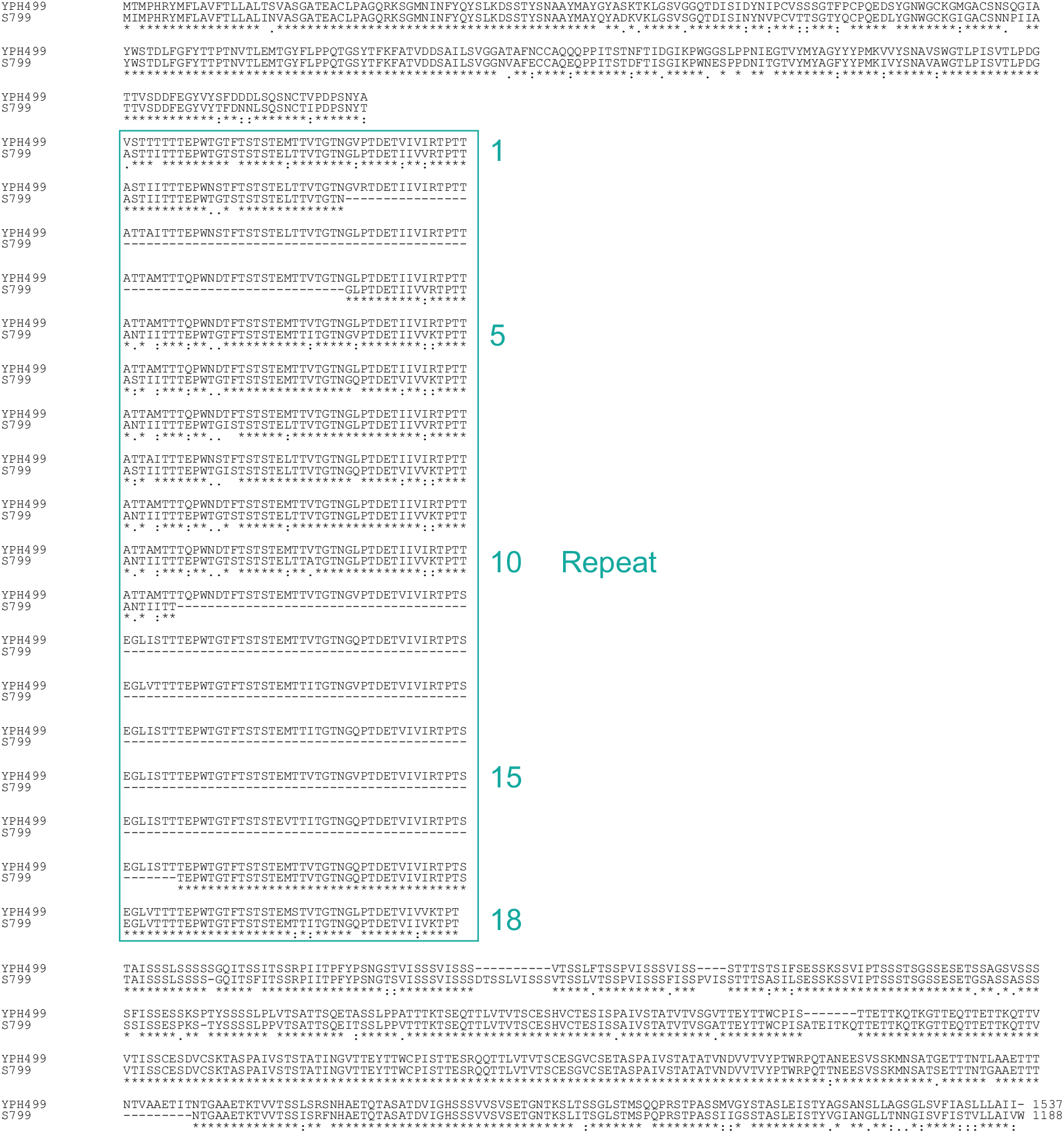
Amino acid sequence alignment of Flo1 protein between YPH499 (upper) and S799 (bottom).

**Supplementary Fig. 4.**
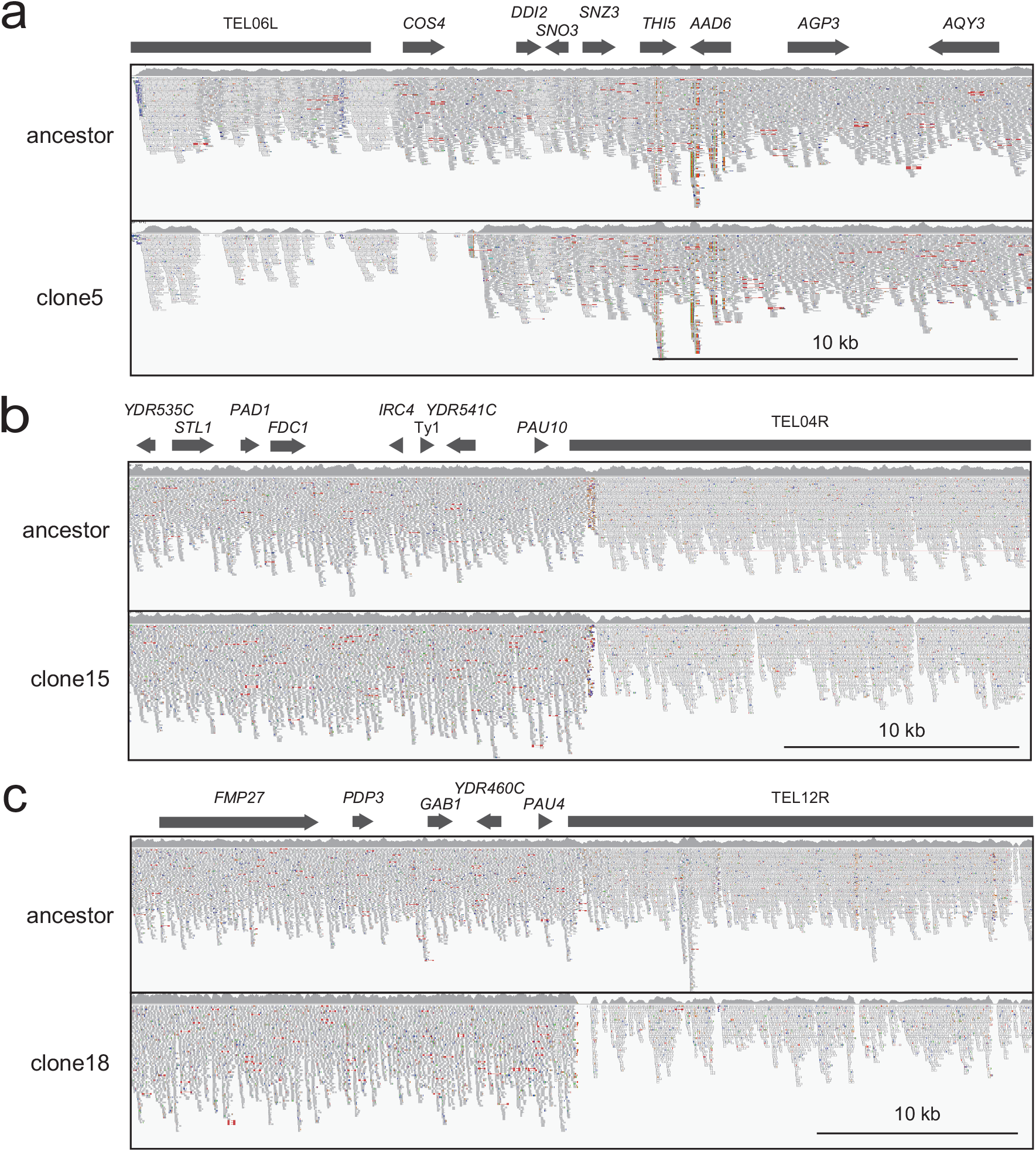
Telomeric deletion in experimentally evolved clones. Sequence-read alignments for the telomeric regions of the (a) left arm of chromosome VI in clone 5, (b) right arm of chromosome IV in clone 15, and (c) right arm of chromosome XII in clone 18 are visualized by IGV.

**Supplementary Fig. 5.**
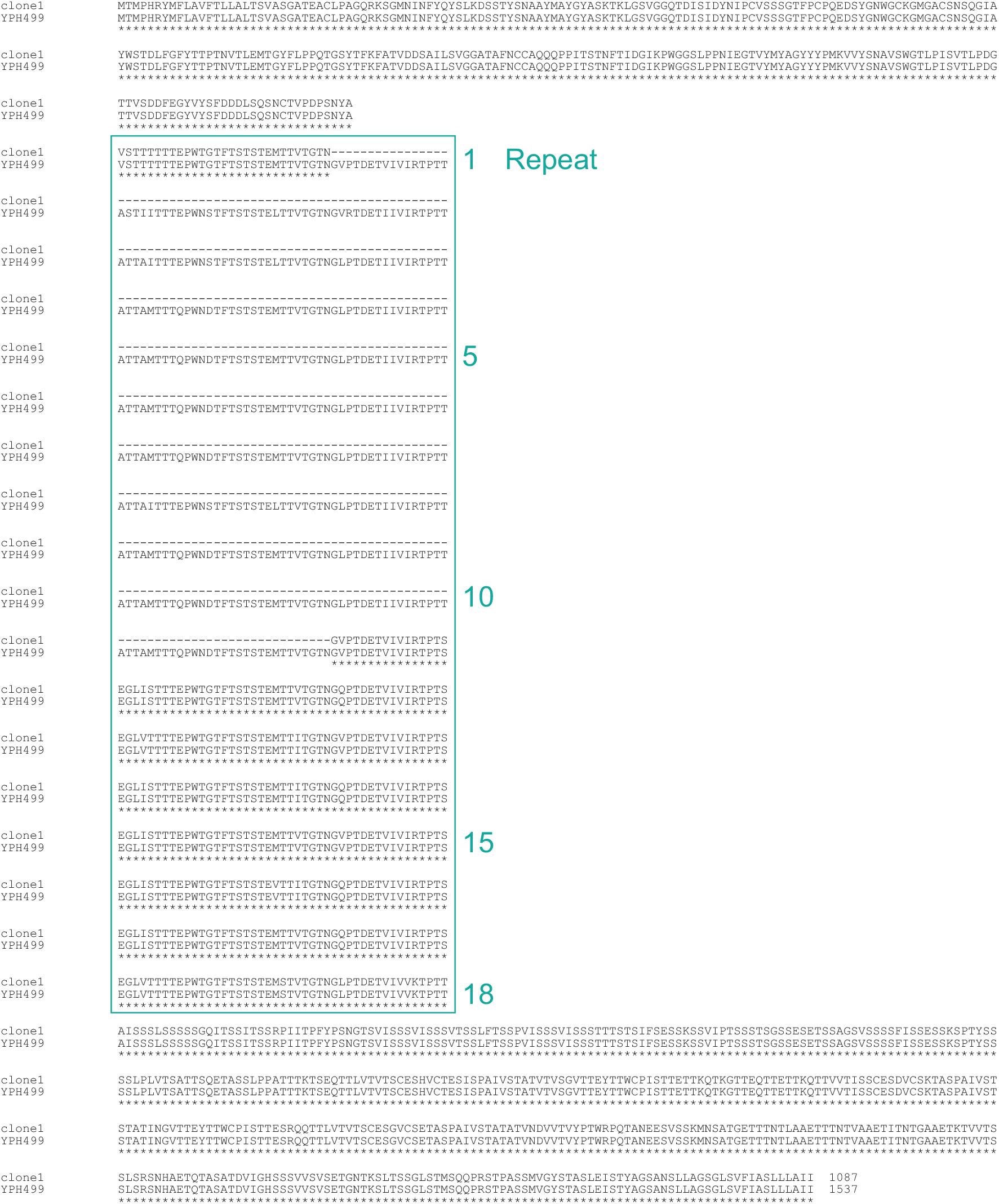
Amino acid sequence alignment of Flo1 protein between experimentally evolved clone 1 (upper) and YPH499 (bottom).

**Supplementary Table 1-1.** Structural variants of TAQed strains.

| Strain | Rearrange type | Chromosome | Position | Mutation |
| --- | --- | --- | --- | --- |
| m126 | SNV | YPH499chr08 | 138,529 | C > G |
| m130 | SNV | S799chr13 | 123,861 | G > A |
| m131 | SNV | YPH499chr08 | 147,194 | C > A |
| m131 | SNV | S799chr15 | 64,350 | A > G |
| m134 | SNV | YPH499chr08 | 70,964 | A > G |
| m134 | SNV | YPH499chr13 | 516,844 | A > C |
| m135 | SNV | YPH499chr04 | 1,517,959 | T > C |
| m144 | SNV | YPH499chr06 | 271,238 | G > T |
| m157 | SNV | YPH499chr15 | 138,794 | T > C |
| m157 | SNV | S799chr07 | 950,059 | G > T |
| m159 | SNV | S799chr12 | 840,323 | G > A |
| m161 | SNV | S799chr07 | 142,504 | A > G |
| m166 | SNV | S799chr13 | 585,981 | C > T |
| m172 | SNV | S799chr01 | 158,942 | T > C |
| m172 | SNV | S799chr05 | 413,007 | G > A |
| m174 | SNV | YPH499chr07 | 17,393 | T > G |
| m177 | SNV | S799chr09 | 286,867 | T > G |
| m179 | SNV | S799chr02 | 503,926 | G > A |
| Strain | Rearrange type | Chromosome | Start | End |
| m126 | SGC | YPH499chr09 | 387,500 | 390,900 |
| m126 | SGC | YPH499chr16 | 165,500 | 167,700 |
| m131 | SGC | YPH499chr08 | 140,500 | 146,400 |
| m135 | SGC | S799chr07 | 624,900 | 632,000 |
| m146 | SGC | S799chr02 | 561,900 | 562,900 |
| m149 | SGC | S799chr02 | 710,200 | 718,500 |
| m159 | SGC | YPH499chr07 | 959,400 | 965,000 |
| m161 | SGC | YPH499chr10 | 151,800 | 161,900 |
| m162 | SGC | YPH499chr05 | 252,600 | 255,000 |
| m163 | SGC | YPH499chr05 | 177,400 | 187,000 |
| m164 | SGC | S799chr15 | 565,802 | 570,600 |
| m165 | SGC | YPH499chr05 | 315,805 | 326,000 |
| m168 | SGC | YPH499chr01 | 164,100 | 194,300 |
| m169 | SGC | YPH499chr01 | 197,300 | 208,100 |
| m170 | SGC | S799chr04 | 728,500 | 729,200 |
| m174 | SGC | YPH499chr04 | 1,033,100 | 1,037,800 |
| m174 | SGC | YPH499chr07 | 17,500 | 30,700 |
| m174 | SGC | YPH499chr13 | 283,700 | 293,600 |
| m174 | SGC | YPH499chr13 | 301,400 | 303,300 |
| m175 | SGC | YPH499chr05 | 308,300 | 318,600 |
| m177 | SGC | YPH499chr15 | 519,100 | 521,100 |
| m178 | SGC | YPH499chr05 | 243,100 | 251,000 |

Blue data represent the approximate rearrangement regions, whereas black data represent precise positions. (SNV; Single Nucleotide Variation, SGC; Short Gene Conversion)

**Supplementary Table 1-2.** Structural variants of TAQed strains.

| Strain | Rearrange type | Left arm | Right arm | Rearranged position |  |
| --- | --- | --- | --- | --- | --- |
| m131 | BIR | S799chr16 | YPH499chr16 | 46,600 |  |
| m134 | BIR | YPH499chr01 | S799chr01 | 204,400 |  |
| m134 | BIR | S799chr13 | YPH499chr13 | 829,000 |  |
| m135 | BIR | S799chr05 | YPH499chr05 | 272,200 |  |
| m135 | BIR | YPH499chr09 | S799chr09 | 287,000 |  |
| m146 | BIR | S799chr05 | YPH499chr05 | 263,900 |  |
| m149 | BIR | YPH499chr02 | S799chr02 | 719,800 |  |
| m149 | BIR | S799chr05 | YPH499chr05 | 251,500 |  |
| m159 | BIR | S799chr07 | YPH499chr07 | 234,700 |  |
| m159 | BIR | S799chr05 | YPH499chr05 | 356,500 |  |
| m161 | BIR | YPH499chr10 | S799chr10 | 151,800 |  |
| m162 | BIR | S799chr05 | YPH499chr05 | 266,700 |  |
| m163 | BIR | S799chr05 | YPH499chr05 | 199,600 |  |
| m165 | BIR | S799chr05 | YPH499chr05 | 339,300 |  |
| m166 | BIR | YPH499chr05 | S799chr05 | 260,400 |  |
| m168 | BIR | YPH499chr01 | S799chr01 | 194,300 |  |
| m170 | BIR | YPH499chr04 | S799chr04 | 752,400 |  |
| m170 | BIR | S799chr15 | YPH499chr15 | 410,800 |  |
| m172 | BIR | YPH499chr01 | S799chr01 | 172,600 |  |
| m172 | BIR | YPH499chr09 | S799chr09 | 326,400 |  |
| m173 | BIR | S799chr05 | YPH499chr05 | 253,000 |  |
| m174 | BIR | S799chr05 | YPH499chr05 | 319,500 |  |
| m175 | BIR | S799chr05 | YPH499chr05 | 330,300 |  |
| m177 | BIR | S799chr15 | YPH499chr15 | 500,000 |  |
| m178 | BIR | S799chr05 | YPH499chr05 | 263,600 |  |
| Strain | Rearrange type | Chromosome | Alteration |  |  |
| m123 | Aneuploidy | YPH499chr01 | Loss |  |  |
| m126 | Aneuploidy | YPH499chr01 | Loss |  |  |
| m130 | Aneuploidy | YPH499chr01 | Loss |  |  |
| m133 | Aneuploidy | YPH499chr01 | Loss |  |  |
| m144 | Aneuploidy | YPH499chr01 | Loss |  |  |
| m150 | Aneuploidy | YPH499chr01 | Loss |  |  |
| m157 | Aneuploidy | YPH499chr01 | Loss |  |  |
| m159 | Aneuploidy | YPH499chr11 | Loss |  |  |
| m160 | Aneuploidy | YPH499chr01 | Loss |  |  |
| m161 | Aneuploidy | YPH499chr01 | Loss |  |  |
| m166 | Aneuploidy | YPH499chr01 | Loss |  |  |
| m166 | Aneuploidy | YPH499chr12 | Gain |  |  |
| m179 | Aneuploidy | YPH499chr01 | Loss |  |  |
| Strain | Rearrange type | Chromosome | Region | Chromosome | Region |
| m131 | TL | YPH499chr01 | 161,900 (Ty element) | YPH499chr16 | 56,900 (Ty element) |
| m159 | Circularization | YPH499chr15 | 26,353 | YPH499chr15 | 466,447 |
| m172 | Deletion | YPH499chr08 | 193,991 | YPH499chr08 | 195,216 |
| m177 | TL | YPH499chr01 | 161,900 (Ty element) | S799chr03 | 178,000 (Ty element) |

Blue data represent the approximate rearrangement regions, whereas black data represent precise positions. (BIR; Break-induced repair, TL; Translocation)

## Notes

### Competing Interest Statement

The authors have declared no competing interest.

## References

1. Swinnen, S., Thevelein, J. M. & Nevoigt, E. Genetic mapping of quantitative phenotypic traits in Saccharomyces cerevisiae. FEMS Yeast Res. 12, 215–227 (2012).

2. Mauricio, R. Mapping quantitative trait loci in plants: uses and caveats for evolutionary biology. Nat. Rev. Genet. 2, 370–381 (2001).

3. Wellcome Trust Case Control Consortium. Genome-wide association study of 14,000 cases of seven common diseases and 3,000 shared controls. Nature 447, 661–678 (2007).

4. Uffelmann, E. et al. Genome-wide association studies. Nat Rev Methods Primers 1, 1–21 (2021).

5. Peter, J. et al. Genome evolution across 1,011 Saccharomyces cerevisiae isolates. Nature 556, 339–344 (2018).

6. Zhu, Z. et al. Integration of summary data from GWAS and eQTL studies predicts complex trait gene targets. Nat. Genet. 48, 481–487 (2016).

7. Wainberg, M. et al. Opportunities and challenges for transcriptome-wide association studies. Nat. Genet. 51, 592–599 (2019).

8. Zeng, B. et al. Multi-ancestry eQTL meta-analysis of human brain identifies candidate causal variants for brain-related traits. Nat. Genet. 54, 161–169 (2022).

9. GTEx Consortium. The GTEx Consortium atlas of genetic regulatory effects across human tissues. Science 369, 1318–1330 (2020).

10. Li, Y. et al. Patterns of somatic structural variation in human cancer genomes. Nature 578, 112–121 (2020).

11. Weischenfeldt, J., Symmons, O., Spitz, F. & Korbel, J. O. Phenotypic impact of genomic structural variation: insights from and for human disease. Nat. Rev. Genet. 14, 125–138 (2013).

12. Cassidy, S. B., Schwartz, S., Miller, J. L. & Driscoll, D. J. Prader-Willi syndrome. Genet Med 14, 10–26 (2012).

13. Meyer-Lindenberg, A., Mervis, C. B. & Berman, K. F. Neural mechanisms in Williams syndrome: a unique window to genetic influences on cognition and behaviour. Nat Rev Neurosci 7, 380–393 (2006).

14. Rowley, J. D. Letter: A new consistent chromosomal abnormality in chronic myelogenous leukaemia identified by quinacrine fluorescence and Giemsa staining. Nature 243, 290–293 (1973).

15. Muramoto, N. et al. Phenotypic diversification by enhanced genome restructuring after induction of multiple DNA double-strand breaks. Nat. Commun. 9, 1995 (2018).

16. Tanaka, H., Muramoto, N., Sugimoto, H., Oda, A. H. & Ohta, K. Extended TAQing system for large-scale plant genome reorganization. Plant J. 103, 2139–2150 (2020).

17. Yasukawa, T. et al. TAQing2.0 for genome reorganization of asexual industrial yeasts by direct protein transfection. Commun Biol 5, 144–13 (2022).

18. Liti, G. et al. Population genomics of domestic and wild yeasts. Nature 458, 337–341 (2009).

19. Kobayashi, O., Hayashi, N., Kuroki, R. & Sone, H. Region of FLO1 proteins responsible for sugar recognition. J. Bacteriol. 180, 6503–6510 (1998).

20. Watari, J. et al. Molecular cloning and analysis of the yeast flocculation gene FLO1. Yeast 10, 211–225 (1994).

21. Verstrepen, K. J., Jansen, A., Lewitter, F. & Fink, G. R. Intragenic tandem repeats generate functional variability. Nat. Genet. 37, 986–990 (2005).

22. Kobayashi, O., Suda, H., Ohtani, T. & Sone, H. Molecular cloning and analysis of the dominant flocculation gene FLO8 from Saccharomyces cerevisiae. Mol. Gen. Genet. 251, 707–715 (1996).

23. Fujita, A. et al. Domains of the SFL1 protein of yeasts are homologous to Myc oncoproteins or yeast heat-shock transcription factor. Gene 85, 321–328 (1989).

24. Pan, X. & Heitman, J. Protein kinase A operates a molecular switch that governs yeast pseudohyphal differentiation. Mol. Cell. Biol 22, 3981–3993 (2002).

25. Liu, H., Styles, C. A. & Fink, G. R. Saccharomyces cerevisiae S288C has a mutation in FLO8, a gene required for filamentous growth. Genetics 144, 967–978 (1996).

26. Song, Q., Johnson, C., Wilson, T. E. & Kumar, A. Pooled segregant sequencing reveals genetic determinants of yeast pseudohyphal growth. PLoS Genet. 10, e1004570 (2014).

27. Hope, E. A. et al. Experimental Evolution Reveals Favored Adaptive Routes to Cell Aggregation in Yeast. Genetics 206, 1153–1167 (2017).

28. Ryan, O. et al. Global gene deletion analysis exploring yeast filamentous growth. Science 337, 1353–1356 (2012).

29. Robertson, L. S. & Fink, G. R. The three yeast A kinases have specific signaling functions in pseudohyphal growth. PNAS 95, 13783–13787 (1998).

30. Reynaud, K., Brothers, M., Ly, M. & Ingolia, N. T. Dynamic post-transcriptional regulation by Mrn1 links cell wall homeostasis to mitochondrial structure and function. PLoS Genet. 17, e1009521 (2021).

31. Rodriguez, M. E., Orozco, H., Cantoral, J. M., Matallana, E. & Aranda, A. Acetyltransferase SAS2 and sirtuin SIR2, respectively, control flocculation and biofilm formation in wine yeast. FEMS Yeast Res. 14, 845–857 (2014).

32. Park, K. C. et al. Purification and characterization of UBP6, a new ubiquitin-specific protease in Saccharomyces cerevisiae. Arch Biochem Biophys 347, 78–84 (1997).

33. Papa, F. R. & Hochstrasser, M. The yeast DOA4 gene encodes a deubiquitinating enzyme related to a product of the human tre-2 oncogene. Nature 366, 313–319 (1993).

34. Yin, Y. & Petes, T. D. Genome-wide high-resolution mapping of UV-induced mitotic recombination events in Saccharomyces cerevisiae. PLoS Genet. 9, e1003894 (2013).

35. Sadhu, M. J., Bloom, J. S., Day, L. & Kruglyak, L. CRISPR-directed mitotic recombination enables genetic mapping without crosses. Science 352, 1113–1116 (2016).

36. Laureau, R. et al. Extensive Recombination of a Yeast Diploid Hybrid through Meiotic Reversion. PLoS Genet. 12, e1005781 (2016).

37. Furuse, M. et al. Distinct roles of two separable in vitro activities of yeast Mre11 in mitotic and meiotic recombination. EMBO J. 17, 6412–6425 (1998).

38. Bahler, J. et al. Heterologous modules for efficient and versatile PCR-based gene targeting in Schizosaccharomyces pombe. Yeast 14, 943–951 (1998).

39. Sato, M., Dhut, S. & Toda, T. New drug-resistant cassettes for gene disruption and epitope tagging in Schizosaccharomyces pombe. Yeast 22, 583–591 (2005).

40. Curran, B. P. & Bugeja, V. C. Protoplast fusion in Saccharomyces cerevisiae. Methods Mol. Biol. 53, 45–49 (1996).

41. Li, H. & Durbin, R. Fast and accurate short read alignment with Burrows-Wheeler transform. Bioinformatics 25, 1754–1760 (2009).

42. Garrison, E. & Marth, G. Haplotype-based variant detection from short-read sequencing. Preprint at http://arxiv.org/abs/1207.3907 (2012).

43. Li, H. A statistical framework for SNP calling, mutation discovery, association mapping and population genetical parameter estimation from sequencing data. Bioinformatics 27, 2987–2993 (2011).

44. Robinson, J. T. et al. Integrative genomics viewer. Nat. Biotechnol. 29, 24–26 (2011).

45. Li, H. et al. The Sequence Alignment/Map format and SAMtools. Bioinformatics 25, 2078–2079 (2009).

46. Hirota, K. et al. Stepwise chromatin remodelling by a cascade of transcription initiation of non-coding RNAs. Nature 456, 130–134 (2008).

47. Galipon, J., Miki, A., Oda, A., Inada, T. & Ohta, K. Stress-induced lncRNAs evade nuclear degradation and enter the translational machinery. Genes Cells 18, 353–368 (2013).

